# Modelling Glioma Stem Cell-mediated Tumorigenesis Using Zebrafish PDX Systems

**DOI:** 10.1101/2024.02.27.580776

**Authors:** Hema Priya Mahendran, Alan Cieslukowski, Dorota Lubanska, Nicholas Philbin, Keith Franklin Stringer, Philip Habashy, Mat Stover, Ana C. deCarvalho, Mohamed Soliman, Abdalla Shamisa, Lisa Porter

## Abstract

Glioblastoma is an aggressive brain tumour associated with high post-therapy recurrence and very poor survival rates. One of the factors contributing to the aggressive nature of this disease is the level of heterogeneity seen at the phenotypic and genetic level. Glioma Stem Cells (GSCs) are stem-like cells within the tumour with the ability to self-renew and give rise to different types of cells within the tumour, hence giving rise to the heterogeneity found in glioblastoma. GSCs are often implicated in the resistance of glioma to standard of care radiation and chemotherapy. The physical niche within a tumour mass supports stemness and aggressive characteristics of GSCs, hence, experimental systems providing a relevant tumour microenvironment are critical for adequate assessment of molecular mechanisms regulating GSC populations. Although, mouse models are a staple of an *in vivo* experimental design, they are neither time-nor cost-efficient. *Danio rerio* (zebrafish) patient-derived xenografts (PDXs) overcome several of the obstacles of the mammalian systems. Zebrafish constitute a high throughput, easily reproducible experimental platform allowing for life relevant investigation into the aggressiveness of GSC populations. This chapter describes methods required for generation of zebrafish PDXs to study aspects of GSC-mediated tumorigenesis and interactions with the tumour microenvironment. Consistency between labs for these experiments is required to move the discovery of effective treatments for glioblastoma moving forward.

## Introduction

Glioblastoma (GBM) or grade IV, IDH-positive, astrocytoma is characterized by poor patient prognosis and limited sensitivity to standard of care treatment^1^. GBM accounts for 50% of all gliomas and 5-year survival rate of less than 10%^2,3^. Glioma stem cells (GSCs), a sub-population of immature neural precursor cells within the GBM mass, resemble normal neural stem cells in their ability of self-renew and differentiate into multiple lineages. GSCs are resistant to conventional therapies and cause tumour recurrence leading to relapse following standard of care therapy^4^. Studying GSCs *in vivo* can aid in the discovery of unique molecular and biological characteristics that can be targeted to improve the treatment of GBM.

In recent years, the role of the tumour microenvironment (TME) in the progression of cancer has become very apparent^8,9^. The TME consists of various structural and cellular components within and around the tumour cells. The most prominent of these components include stromal cells such as fibroblasts, pericytes, endothelial cells, and various cells of the immune system. Studies have shown that cancer can co-opt these cells and use their various biological roles to progress tumour growth^9^. Thus, it is of critical importance to dissect the molecular mechanisms by which the cells and molecules of the TME are able to impact tumour evolution. Such work has the potential to reveal vulnerabilities of cancer that can be targeted for improved therapeutic outcome.

The value of animal model systems for the study of TME in glioma biology cannot be under emphasized. Decades of study have revealed conservation of important aspects of the TME in a host of laboratory animal models. Human cancer biology can further benefit from these models by combining human cells either orthotopically (into the tissues where they are found in the patient) or heterotopically (into sites with a TME similar to the site of origin of the tumour). These models are referred to as patient-derived xenograft (PDX) models and they permit study of how the cells from human patients respond when exposed to the system of an intact organism. Mouse PDX models are the most common and are valuable in providing several key components of the TME necessary to maintain the original tumour cytoarchitecture^10^. Despite their advantages, injecting human cells into mouse models requires the use of immunocompromised animals and hence, mouse-PDX models lack crucial components of the immune system. They are also time consuming and expensive.

Zebrafish embryo PDXs on the other hand, constitute a high throughput and relatively inexpensive model for tumour growth and drug response. Zebrafish PDX models highly predictive of clinical response to chemotherapeutic drugs^11^. Clinical relevance of the PDX models is validated to be as high as 90%^12^. Zebrafish embryos have an additional benefit in that they possess an active innate immune system at the time of injection of the human cells but lack the presence of a functional adaptive immune system until embryonic day 21, which permits successful engraftment of human cells^13^. The use of the zebrafish strain Casper, lacking melanocytes responsible for pigmentation, renders these fish nearly transparent and provides the added benefit of improving detection of fluorescent signals that can be incorporated into the xenografted cells prior to injection. All of these benefits make the PDX-Zebrafish model particularly attractive for the study of how human cells interact with the TME to drive aspects of cancer biology including tumour progression and metastasis. The genetic pliability of the model further allows experimental manipulation of TME components to make conclusions with regard to the causality of specific molecules and/or cell types within the TME^14^.

In this chapter, we describe the labelling of GSCs for tracking of the cells in PDX-zebrafish systems over time. We outline the process of microinjection of GSCs into zebrafish to generate PDXs that are reflective of GBM behaviour in patients, and we cover protocols for the analysis of tumour burden, migration and recruitment of GSCs to the TME of GBM. We also provide instructions for the staining of zebrafish with haematoxylin & eosin and immunohistochemistry of zebrafish sections to investigate tissue architecture and study the expression and localization of specific proteins. Lastly, the detailed protocols outlined can be applied to the study of the interaction of GSCs and various cell types and components of the TME, particularly those that have been shown to be of most importance in GBM. In this chapter we also cover various transgenic zebrafish models that allow for the identification and tracking of TME interactions with human cells. The transgenic zebrafish models listed are excellent systems to study GSC biology in the context of the TME.

Collectively, this chapter aims to provide researchers with insight into a high throughput and cost-efficient *in vivo* model that has been shown to be highly reflective of the behaviour of GBM seen in patients^12^. The sharing of detailed protocols will support consistency of results that have the potential to support novel discovery and advance of new treatments to improve outcomes in the clinical treatment of GBM.

## Materials

### Cell Lines

1. Patient-derived glioblastoma cell lines (GSCs) were obtained from the Dr. Lisa A. Porter lab brain tumour clinical trial partnership with Windsor Regional Hospital in Windsor, Ontario, Canada and Henry Ford Hospital in Detroit, Michigan, United States of America. Specifically, the line used for the methods outlined here are HF3077 and HF3253 (See Note #1).
2. U251-MG cells were kindly provided by (Dr. Rutka, The Hospital for Sick Children, Toronto, Ontario, Canada), but can be obtained from American Culture Type Collection (ATCC; CA#CRL-1730).

### Cell Culture Reagents

#### GSC culture

GSC media formulation:

1. Dulbecco’s Modified Eagle’s Medium/Hams F-12 50/50 mix (DMEM/F-12 50/50) (Corning, Sigma; CA#45000-350)
2. Epidermal Growth Factor (EGF) (Gibco, ThermoFisher; CA#PHG0311)
3. Human Basic Fibroblast Growth Factor (hBFGF) (Sigma; CA#FO291)
4. 10mg/mL Gentamicin (Gibco, ThermoFisher; CA#15710064)
5. 100X N-2 Supplement (Gibco, ThermoFisher; CA17502048)
6. 100X Antibiotic-Antimycotic (Gibco, ThermoFisher; CA#15240-062)
7. Bovine Serum Albumin (BSA) (Sigma; CA#A4919-5G)

#### GSC culture supplies

8. pH 7.4 PBS (Gibco, ThermoFisher; CA# 10010023)

9. T25 Red Cap Tissue Culture Flask (Sarstedt; CA#83.3910.002)

10. 15mL, blue screw cap, conical bottom tube (Grenier; CA#188261)

#### GSC Infection Media

1. GSC media minus antibiotics and minus BSA

#### U251-MG Cells

1. 0.25% Trypsin (Hyclone; #SH3023601)
2. pH 7.4 1X PBS
3. 15mL, blue screw cap, conical bottom tube (Grenier; CA#188261)

#### U251-MG Media

1. Minimum Essential Medium Eagle (EMEM) (Sigma; CA#10128-608)
2. 10% Fetal Bovine Serum (FBS) (Gibco; CA#10437028)
3. 1X Non-Essential Amino Acids (NEAA) (Sigma; CA#M7145)
4. Puromycin (Sigma; CA#P8833)

#### U251-MG Infection Media

1. Minimum Essential Medium Eagle (EMEM) (Sigma; CA#10128-608)
2. 1X Non-Essential Amino Acids (NEAA) (Sigma; CA#M7145)

#### Transgenic Zebrafish Model

1. The Tg(fli1a:dsRED) transgenic zebrafish model was provided by the Dalhousie University Zebrafish Core Facility in Halifax, Nova Scotia, Canada.

#### Injection of GBM Cells in Zebrafish

1. 60X E3 Media: 286.3mg NaCl, 12.5mg KCl, 50mg CaCl_2_, 81.5mg MgSO4, 1L H_2_O, 20uL 0.3% methylene blue (synthesized at University of Windsor, Ontario, Canada)
2. pLBS-mScarlet (Addgene; CA#129337)
3. pLEX304mNeonGreen (Addgene; CA#162034)
4. Vybrant DiD Cell-Labelling Solution (ThermoFisher; CA#V22887)
5. Vybrant DiO Cell-Labelling Solution (ThermoFisher; CA#V22886)
6. pH 7.0 15mM Tricaine (Sigma; CA#A-5040)
7. Nanoject III Injector (Drummond Scientific Company; CA#3-000-207)

#### Tissue Processing

1. Precleaned Superfrost Plus Microscope Slides (Fisherbrand; CA#12-550-15)
2. 4% PFA in PBS (Santa Cruz Biotechnology; CA#SC281-692)
3. pH 7.0 15mM Tricaine (Sigma; CA#A-5040)
4. pH 7.0 0.35M EDTA
5. 100% EtOH
6. 95% EtOH
7. 80% EtOH
8. 70% EtOH
9. Xylenes (ACP Chemicals; CA#X-0250)
10. Paraplast Plus (McCormick Scientific; CA#502004)
11. Low Melting Point Agarose (Sigma-Aldrich; CA#AG9045-5G)
12. Microscope Cover Glass (UltiDent; CA#170-C2260)
13. Super Pap Pen Liquid Blocker for IHC (Mercedes Scientific; CA#SPM0928)
14. pH 7.4 1X PBS.
15. DIH2O
16. Parafilm (Fisher Scientific; CA#13-374-10)
17. Metal Rack
18. Slide Staining Jars
19. Permount Mounting Solution (Fisher Chemical; CA#SP15-100)
20. Hematoxylin Gill’s Formula (Vector Laboratories, Inc; CA#H-3401)
21. Eosin Y
22. VECTASHIELD (Vector Laboratories, Inc.; CA#H-1000)
23. Citrate Buffer: 2.93g Na3C6H5O7, pH to 6.0, 500uL Tween20
24. Blocking Solution: 3% Albumin Fraction V (Bio Basic; CA#9048-46-8), 0.1% Tween20 (ACP Chemicals; CA#T-9780) in 1X PBS
25. DAPI Solution (BD Pharmingen; CA#564907)
26. FITC Anti-Human CD274 (B7-H1, PD-L1) Antibody (BioLegend; CA#393606)
27. Lymphocyte Cytosolic Protein (LCP) (GeneTex; CA#GTX124420)
28. Goat anti-Mouse IgG (H+L) Cross-Adsorbed Secondary Antibody, Texas Red-X (ThermoFisher; CA#T-6390)
29. Goat Anti-Rabbit IgG H&L (Cy5) Preadsorbed (Abcam; CA#ab6564)

## Methods

### Cell Culture

#### GSCs

1. Subculture GSCs by allowing neurospheres formed by the cells to sediment in a 15mL conical tube.
2. Aspirate and remove the media.
3. Add 2mL of pH 7.4 PBS.
4. After 10 minutes of exposure to pH 7.4 PBS, gently disrupt the neurospheres with a P1000 pipette. Repeat this step every 10 minutes until a single cell suspension is formed.
5. Centrifuge the single cell suspension at 800g for 5 minutes.
6. Aspirate and remove the pH 7.4 PBS.
7. Resuspend in 1mL of warm growth media and add the suspension to 10mL of warm growth media in a T25 red cap tissue culture flask.
8. Place the cells in an incubator set at 37 degrees Celsius and 5% CO_2_.

#### U251-MG Cells

1. Aspirate and remove all the growth media from the plate.
2. Wash the plate with 1mL of pH 7.4 PBS.
3. Aspirate and remove the 1mL of pH 7.4 PBS.
4. Add 1mL of 0.25% trypsin.
5. Incubate the plate for 5 minutes at 37 degrees Celsius and 5% CO_2_.
6. After 5 minutes, remove the plate from the incubator and use 1mL of growth media to wash the cells off of the plate and collect the cells into a 15mL conical tube.
7. Centrifuge the cells at 1000RPM for 5 minutes.
8. Aspirate and remove the supernatant and resuspend the cells in 1mL of warm growth media.
9. Add 5-10uL of the cell suspension to 10mL of warm growth media in a 10cm plate.

### Vybrant Cell-Labelling Solution Staining (See Note #2)

1. Collect GSCs or U251-MG cells as a single cell suspension in a 15 mL conical tube with growth media (see protocol(s) above).
2. Determine the live cell count of the single cell suspension using a haemocytometer.
3. Resuspend 1x10^6^ cells in 1mL of HBTC+ media or U251-MG media and add 10uL of either Vybrant DiO or Vybrant DiI Cell-Labelling Solution (See Note #3).
4. Incubate the sample at 37 degrees Celsius and 5% CO_2_ for 40 minutes.
5. Touch vortex the sample every 10 minutes.
6. After 40 minutes, centrifuge the sample (at 800g for GSCs or 1000RPM for U251-MG cells) for 5 minutes.
7. Aspirate and remove the supernatant.
8. Resuspend the sample in 100 uL of clean DMEM-F-12 media and store on ice until injection (See Note #4).

### Lentiviral Transduction (See Note #5)

1. Collect GSCs or U251-MG cells as a single cell suspension in a 15 mL conical tube with growth media (see protocol(s) above).
2. Determine the live cell count of the single cell suspension using a haemocytometer.
3. Seed the cells at a density of 1 x 10^4^ cells/well and 100uL of HBTC+ infection media or U251-MG infection media/well.
4. Add 2uL of lentivirus containing pLBS-mScarlet or pLEX304mNeonGreen to each well.
5. Place the cells into the incubator overnight at 37°C and 5% CO_2_.
6. The next day, remove the HBTC+ or U251-MG infection media and replace with 100uL of HBTC+ or U251-Mg media.
7. Allow the cells to adequately recover. Monitor by observing indicators of healthy cells under the microscope.
8. Once the cells have recovered and reach confluency, subculture according to the protocol(s) mentioned above to expand the populations.

### FACS of Cells Infected With pLBS-mScarlet or pLEX304mNeonGreen

1. Collect GSCs or U251-MG cells infected with pLBS-mScarlet or pLEX304mNeonGreen as a single cell suspension in a 15 mL conical tube with growth media (see protocol(s) above).
2. Determine the live cell count of the single cell suspension using a haemocytometer.
3. Resuspend 1x10^6^ cells in 1mL of 2mM PBS EDTA (See Note #6).
4. Strain the cells through a 70um cell strainer prior to FACS.

### Agar Plate Preparation

1. For injection, place Zebrafish embryos place on a semi-solid medium made of 0.8% agarose in water.
2. To make the agarose plates, fill the lid of the petri dish with approximately 50 mL of agarose to fill the entire area.
3. Allow the plates to reach room temperature before use for zebrafish embryo injections.

### Zebrafish GSC Xenografts

1. Tip of the glass needle should be cut open diagonally at a 45° angle at ¾ cm from the tip to yield a sharper tip with a diameter of 5-10µM (See Note #7).
2. The needle should then be filled with the mineral oil through syringe to check the consistent flow.
3. Position the plunger on injector upright and put the needle around and tighten. Make sure there are no air bubbles, and the plunger is all the way up.
4. The oil in the needle can be emptied later until only a small volume is left.
5. Zebrafish embryos at 72hrs post fertilization (hpf) can be used to establish xenografts.
6. The embryos can be anaesthetized in 1mL of Tricaine at 40µg/mL concentration in E3 media for approximately 3-5 minutes (See Note #8).
7. On a clean parafilm pipette ∼20µL cell suspension and use it to fill in the needle.
8. Set the injector to 9.5nL volume of injections at the rate of 1µL/Sec (See Note #9).
9. Cells can be either injected on yolk sac or orthotopically into the brain (See Note #10).
10. For yolk sac injection the needle tip should be placed directly above the yolk sac, pierced into centre at 45° angle and inject the cells (See Note #11).
11. For orthotopic brain injection, the needle tip should be directed into the middle of the brain which corresponds to the superficial area above the optic tectum.
12. After injection the embryos should be placed in the regular E3 media for recovery.
13. The embryos can be screened for successful engraftment using Leica EZ4 fluorescence stereo microscope.

### Imaging of Zebrafish PDXs

1. Three to five PDX zebrafish embryos were transferred to agarose plates using Pasteur pipette for imaging. All the embryos were aligned in a similar straight manner and covered with a drop of E3 media to prevent them from drying. Important to note that with too much E3 media, it’s hard to image as the fishes keep moving.
2. The injected embryos are imaged using Leica fluorescent microscope with LAS X software at the indicated timepoints (See Note #12).
3. The images are exported from individual channels and overlay for further analysis.

### Image Analysis-Integrated Density Values Score

1. The images taken at different time points are used to measure the tumour burden changes and migration over time.
2. The images were analysed using ImageJ software.
3. To measure the tumour burden, the fluorescent channel image was imported to the software and converted to 8 bits (black & white).
4. The threshold was set in a way the background fluorescence was eliminated.
5. Then the total number of fluorescent cells was measured as integrated density values and used to analyse tumour burden.
6. Using the same method change in tumour burden at different time points were measured.

### Image Analysis-Tumour Foci

1. Number of tumour foci in each fish was also measured using the similar technique at various sites.
2. For accurate foci count, the fluorescent channel image with scale bar was used.
3. Using the exact length of scalebar in the image, a scale for analysis was set to calculate accurate foci area.
4. The area of interest is selected using one of the shape tools. The dimensions (width & height) of the shape is saved by going to Analyze: Tools: ROI Manager: Add t.
5. Using preview point selection, noise tolerance is adjusted as appropriate, so that only the tumour foci are being counted.
6. Select Output type: Count – this will generate a separate results box.
7. Foci counts in the results box can be copied and pasted into Excel or GraphPad Prism to analyze data.

### Image Analysis-Migration

1. To analyse the migration, ImageJ software was used to measure the migration pattern, distance travelled and directionality of tumour cells within the zebrafish embryo.
2. Multipoint tool is used to select the injection site and X & Y coordinates of the injection site is measured.
3. The measurements were set to centroid under Analyze: Set Measurements and Analyze particles was selected from the toolbar.
4. This will provide the coordinates of each cell metastasized.
5. The farthest moved cell and total distance covered by the individual cells from site of injection was estimated by correcting it with the site of injection.
6. The straightness or directional persistence of cell movement was measured.
7. The measurements from tumour burden and migration were used to generate graphical representations and comparisons between experimental groups.

### Sectioning and Embedding

1. After euthanasia with a 1:25 dilution of 15mM cold tricaine in 1X PBS for 5-10 minutes, collect zebrafish in 4% PFA in a flat-bottom vessel and allow fixation to occur for 24 hours (See Note #13 & Note #14).
2. Then, place the zebrafish in pH 7.0 0.35M EDTA at 21°C to remove calcium and mineralized salts. This will also soften the zebrafish for sectioning.
3. Orient and embed the zebrafish in 1% low melting point agarose after decalcification (See Note #15).
4. Dehydrate the tissue according to the following steps: Incubate in 70% EtOH overnight, 80% EtOH for 45 minutes at 25°C, 95% EtOH for 45 minutes at 25°C, 95% EtOH for 1 hour at 25°C, 100% EtOH for 1 hour at 25°C (repeat three times), xylene for 1 hour at 60°C (repeat two times), paraffin for 1.5 hours at 60°C (repeat two times) and paraffin for 2 hours at 60°C (See Note #16)
5. Take paraffin-embedded blocks and soak in a beaker of ice water for 1 hour in the fridge (See Note #17).
6. Fill water bath and set to 45°C.
7. Once the block is on the stand, set the section diameter to 10um and trim the block until the tissue is present and exposed on the surface.
8. Set the section diameter to 5um and begin sectioning.
9. Once a ribbon of sections is produced, place the ribbon in the 45°C water bath using a pick tool and paintbrush, so the sections do not collapse onto each other.
10. Acquire a Pre-Cleaned Superfrost Microscope Slide and scoop the tissue ribbon from underneath.
11. Leave the slide in a drying rack and then place in a slide box when dry.

### Hematoxylin & Eosin Staining (See Note #18)

1. Bake the slide(s) containing the sections in a metal rack at 65°C for 20 minutes.
2. In the meantime, set up 8 staining jars with one for each of the following unless otherwise specified: xylene (2 staining jars), 100% EtOH (2 staining jars), 95% EtOH, 70% EtOH, DIH_2_O, 1X PBS.
3. In addition, set up a section of parafilm on the bench that will be large enough to hold the slide(s) during staining with hematoxylin and eosin. Tape this section of parafilm down securely.
4. After 20 minutes, remove the slide(s) from the oven and metal rack. Allow the slide(s) to adjust to room temperature for 10 seconds.
5. Using forceps (for all steps involving transfer of the slide(s)), place the slide(s) into the first staining jar containing xylene for 10 minutes. Repeat this step one more time by moving the slide(s) into the second staining jar of xylene.
6. After the second wash in xylene, move the slide(s) into the staining jar with 100% EtOH for 5 minutes. Repeat this step one more time by moving the slide(s) into the second staining jar of 100% EtOH.
7. After the second wash in 100% EtOH, move the slide(s) into the staining jar with 95% EtOH for 2 minutes.
8. After 2 minutes, move the slide(s) into the staining jar with 70% EtOH for 2 minutes.
9. After 2 minutes, briefly place the slide(s) in the staining jar with DIH_2_O before moving the slide(s) specimen side up onto the parafilm set up on the bench.
10. Using the PAP Pen, outline the specimens on the slide to minimize the volume of hematoxylin and eosin required for staining and to ensure the hematoxylin and eosin will cover the specimen adequately.
11. Add 100uL of a 1:4 dilution of hematoxylin in DIH_2_O to the region of the slide(s) surrounded by the PAP Pen. Leave the hematoxylin on for 2 minutes (See Note #19).
12. After 2 minutes, cautiously remove the hematoxylin from the slide(s) with a P200 pipette. Make sure that the specimens remain on the slide. Move the slide(s) into the staining jar with 1X PBS for 2 minutes.
13. After 2 minutes, briefly submerge the slide(s) in DIH_2_O and then place the slide(s) onto the parafilm on the bench.
14. Add 100uL of eosin to the region of slide(s) surrounded by the PAP Pen. Leave the eosin on for 1 minute (See Note #19).
15. Set up 3 more staining jars: 1 with fresh 95% EtOH and 2 with fresh 100% EtOH.
16. After 1 minute, cautiously remove the eosin from the slide(s) with a P200 pipette.
17. Place the slide(s) into the staining jar with fresh 95% EtOH for 5 minutes.
18. After 5 minutes, move the slide(s) into the staining jar with fresh 100% EtOH for 5 minutes. Repeat this step one more time by moving the slide(s) into the second staining jar of fresh 100% EtOH.
19. After the second wash in fresh 100% EtOH, place the slide(s) into the first staining jar with xylene for 5 minutes. For this step, it is possible to reuse the xylene in the first initial staining jar (See Note #20). Repeat this step one more time by moving the slide(s) into the second initial staining jar of xylene.
20. After the second wash in xylene, move the slide(s) onto the parafilm on the bench. Immediately place 1 drop of Permount onto the specimens and place coverslips over the specimens.
21. Leave the slide(s) overnight in a fume hood protected from light.

### Immunohistochemistry

1. Submerge slides in Xylene for 15 minutes, 3 times. (Move from 1 to 2 to 3).
2. Move to 100% EtOH for 5 minutes, 2 times. (Move from 1 to 2).
3. Move to 90-95% EtOH for 5 minutes, then onto 80% EtOH for 5 minutes and finally onto 70% EtOH for 5 minutes.
4. Move to DIH2O and transfer slides from EtOH, submerge for 5 minutes.
5. Microwave the slides in citrate buffer (pH 6.0) for 10 minutes; fill with more citrate buffer or add H2O and microwave another 6 minutes. Let cool to room temperature for 20 minutes.
6. Submerge in water for 7.5 minutes, 2 times (Move from 1 to 2). Total 15 minutes in water. Transfer in 1X PBS to bench. During waiting, fill slide incubator with water.
7. Carefully blot dry around the tissue without touching the mounted tissue spot. Use PAP Pen to mark around the tissue section (See Note #21).
8. Incubate the slides in blocking solution for 20 minutes at room temperature. Add blocking solution onto spot with mounted tissue only with a pipette.
9. After 20 minutes, blot excess serum from sides.
10. Add primary antibody solution (at concentration recommended by the manufacturer) to demarcated sections.
11. Incubate overnight at 4°C in water-filled slide box to keep the inside of the box moist (See Note #22).
12. Following overnight incubation at 4°C, wash slides with room temperature 1X PBS. Perform this step 3 times, with 5-minute washes each time.
13. Add secondary antibody solution (with DAPI solution added) to demarcated sections.
14. Incubate slides for 20 minutes at room temperature.
15. Wash slides with room temperature 1X PBS. Perform this step 3 times, with 5-minute washes each time.
16. Mount slides with one drop of VECTASHIELD, then carefully add slide.
17. Store slides overnight in the dark at room temperature (See Note #23).

### Notes

1. Patient derived GBM lines are cultured for at least ten passages as neurospheres to assure the selection of GSCs which are then validated upon xenografts to nude mice, if they can initiate tumor.
2. If only a membrane-labelling dye is used to label the cells, the fluorescent signal will diminish over time with each cell division/generation. Manufacturers state that the fluorescent signal will be significant for up to 7 generations. This may lead to inaccurate data when studying tumour burden as the cells may be proliferating and increasing in number, but this will be reflected as a decrease in the fluorescence signal observed, resulting in the incorrect conclusion that tumour burden is decreasing over time. For this reason, it is ideal to infect cells with a lentiviral construct encoding a very fluorescent protein such as mNeonGreen or mScarlet for constitutive expression.
3. For the ThermoFisher Vybrant Cell-Labelling Solutions, the recommended staining concentration in the protocol is 1uL of dye/1 x 10^5^ cells/100uL of media. Therefore, depending on the desired number of cells to be stained, the volume(s) required will vary. Always consult the specifications sheet, especially if any membrane-labelling/cyanine dyes from other manufacturers are to be used.
4. The preparation of cells in terms of concentration is mentioned in the last step of the Vybrant Cell-Labelling Solution Staining protocol, however, this step must also be completed prior to injection of GSCs infected with pLBS-mScarlet or pLEX304mNeonGreen and sorted via FACS. Prior to injection, cells should always be resuspended in clean DMEM/F-12 media at a concentration of 1 x 10^6^ cells/100uL media and stored on ice. Cells can be stained with Vybrant Cell-Labelling Solution just prior to this step. However, it is likely that GSCs will be cultured and placed in the incubator after lentiviral infection and FACS, therefore, when using GSCs follow the subculture protocol outlined for these cells and then resuspend in clean DMEM/F-12 media at a concentration of 1 x 10^6^ cells/100uL media and place on ice for injection.
5. For FACS of populations infected with lentivirus in order to obtain pure populations, cells should be resuspended in 2mM PBS EDTA prior to FACS at a concentration of 1 x 10^6^ cells/1mL PBS EDTA. Thus, depending on the desired number of cells to be sorted, the volume of PBS EDTA will need to be adjusted.
6. Although the efficiency of lentiviral infection can be adequate without the use of polybrene, polybrene will greatly enhance the efficiency of lentiviral infection if used in the infection media, leading to more pure populations that can then be injected and provide more accurate results.
7. The glass needles used for zebrafish injections might vary in size and should be optimized based on various factors like volume of cells to be injected, rate of injection, type of needle, plunger etc. Making sharper tip of needle ensures better injections and prevents more tissue damage in embryos. For precise delivery of cells, the microinjection needles are trimmed at the tip at a 45-degree angle.
8. The cells are aspirated to the needle while the zebrafish embryos are being anesthetized. This will prevent the cells from getting sedimented to the bottom of the needle as it is directed downwards and ensures the uniform volume of cells being injected.
9. The injection volume and rate of injection was set based on the zebrafish embryo stage & kept constant throughout the injection. Number of cells to be injected should be optimized based on the rate of proliferation and the size of needle. Excessive injection volumes can result in tissue damage or poor survival.
10. If the injections take longer, there is a risk of embryos drying. Orthotopic brain injections might take longer than yolk sac injections. To prevent the risk of drying, a small volume of E3 media can be added around the fish in agarose plate just enough for the embryos to survive.
11. Identifying the correct injection sites based on the experimental requirements is very important. Common site of injection is yolk sac for embryos and cells can be injected orthotopically at different sites. Other alternative areas of injection include caudal vein, duct of Cuvier, hindbrain, tail muscle, etc (Figure 1A). 72hpf (immediately after hatching) would be an ideal time for injection in yolk sac as it is bigger and easier to inject. Practicing the injections at specific sites assures reproducibility. Important to make sure that cells are not injected in duct of Cuvier unless breast cancer cells, as it causes immediate migration of cells. This affects the results if the experiment is to measure the directionality and movement of cells.
12. Time points for the experiments can vary depending on the type of cells used, their rate of proliferation, number of cells injected and the experiment. For time point experiments, it is better to keep the magnification lower initially as the fish grows and the magnification should be kept consistent between the timepoints. Image settings like magnification and exposure can be saved to keep it constant.
13. Zebrafish must be fixed in a flat-bottom tube/container/vessel in order to achieve equal fixation across all regions of the zebrafish. Do not fix zebrafish in a conical tube as certain regions may not be adequately fixed.
14. Over fixation can result in masking of the epitope so it is paramount to fix the zebrafish for an optimal amount of time and in the optimal fixative.
15. It is important to use low melting point agarose as it ensures that when the zebrafish are later embedded into paraffin wax, the agarose does not melt and the zebrafish orientation is not disturbed.
16. This series of steps is adapted from Copper *et al*^14^. More information about this part of the protocol can be found at the following link: https://bio-atlas.psu.edu.
17. This step allows the blocks to soften resulting in optimal ribbon production of the fish sections.
18. For hematoxylin and eosin staining, the pH of eosin should be 5.0 for proper staining. Check the pH at the beginning of the protocol and if the pH is not 5.0, then use 10 M NaOH and or acetic acid to alter the pH.
19. Hematoxylin and eosin are photosensitive, so it is essential to protect these reagents from light when in use for optimal staining.
20. Whereas the ethanol should be replaced after the initial washes, the xylene from the initial washes can be reused as long as paraffin deposits are not visible at the bottom of the staining jar prior to use for the second washes with xylene.
21. Do not let the tissue section dry during this step. Add a drop of 1X PBS to the spot with the mounted tissue using a pipette after marking with PAP Pen.
22. Depending on the primary antibody utilized and its specifications as per the manufacturer, incubation can be overnight at 4°C or for 1 hour at room temperature. In addition, for primary antibodies that require incubation at 37°C, an Immunohistochemistry Humidity Chamber will need to be set up and the slides will need to be placed into the chamber with primary antibody solution once it has reached 37°C. Make sure to check the slides every 10 minutes for any significant evaporation of the primary antibody solution as drying out will result in the failure of the protocol. Add additional primary antibody solution if needed. Non-primary, secondary treated sections are recommended as staining controls.
23. For long-term storage, wrap slides in aluminium foil and store at 4°C.
24. Zebrafish embryos survive without food for up to 7 days as they utilize the nutrients from yolk sac. With increase in time after hatching, there will be a decrease in size of yolk sac and by day 7, the yolk sac disappears. At this stage the zebrafish should be fed with food or supplied with nutrients in water. If the experiments are longer than 7 days, this might affect the results.
25. The cells that are injected should be high in percentage of viability. Ideally 80-90% or more cell viability is required for good PDX. While the neurospheres are collected, they should be left to dissociate with time and occasional mixing in PBS, instead of pipetting and causing mechanical stress, which affects the cell viability.

## Results

### Xenografts with individual or mixed populations of GSCs

HF3077 and HF3160 GSCs were xenografted into zebrafish yolk sac or brain of 72hpf larvae and analysed as described in Fig 1A. Individual suspension of HF3077 and HF3160 GSCs were injected into the yolk sac (Fig. 1B). Mixed suspension of GSCs HF3077 and HF3160 was grafted. At 24hpi, co-injected cell lines exhibit clear individual signal (Fig. 1C). HF3077 cells represent proneural GBM subtype and exhibit higher proliferation *in vivo* in comparison to mesenchymal HF3160 cells (Fig. 1D). At 5dpi 9.5nl of 10^6^ HF3077 cells increase the number of tumour foci which is correlated with tumour burden further supporting proliferative characteristics of proneural GBM subtype (Fig. 1E).

**Figure 1.**
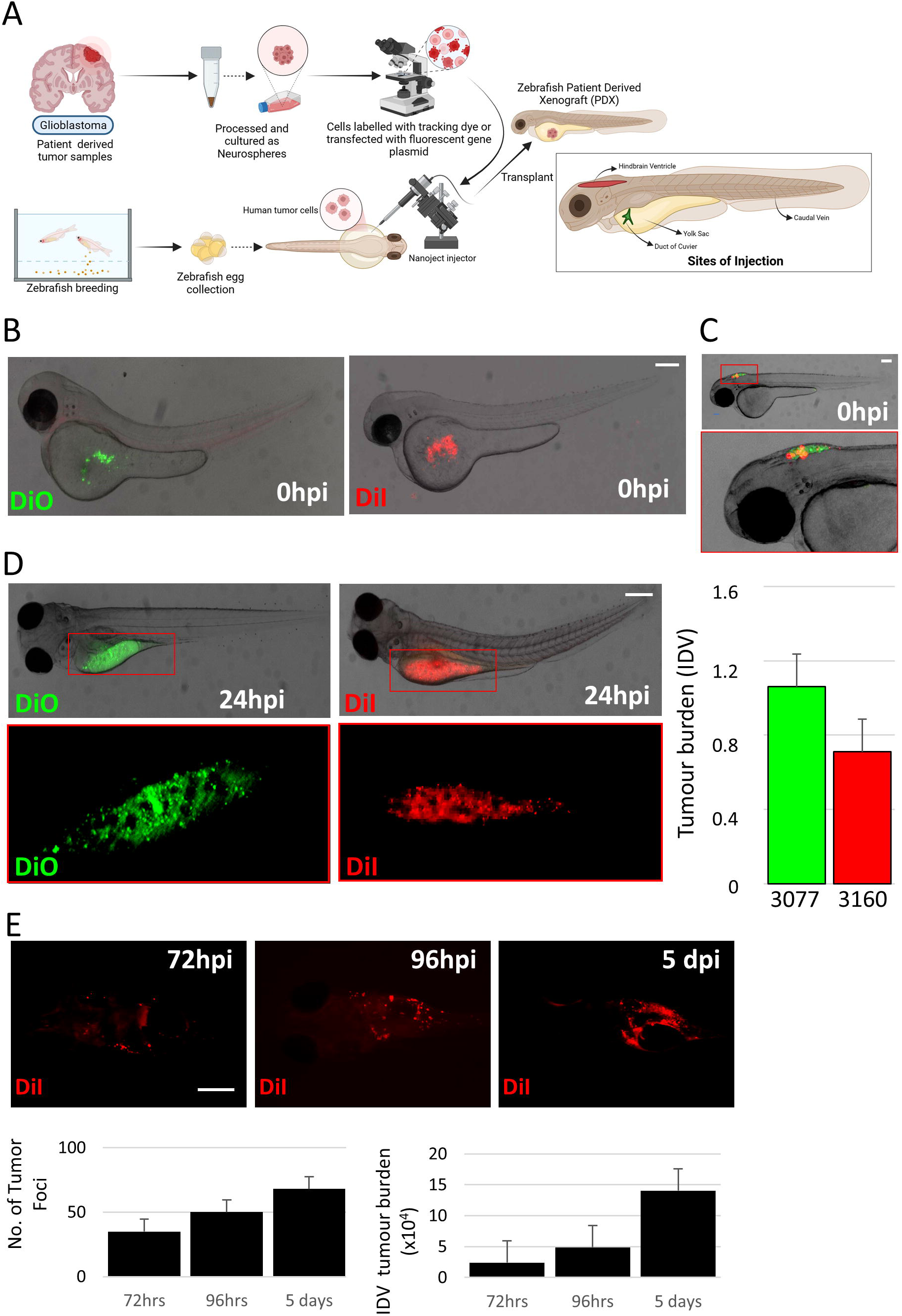
Zebrafish patient derived xenografts (PDX) of GSCs. (A) Schema of the zebrafish PDX workflow. Created with BioRender.com. (B) Zebrafish at 72hpf, injected into yolk sac (0 hours post injection-0hpi) with GSCs HF3077 labelled with DiO (left) and GSCs HF3160 labelled with DiI (right). Micrometer bar: 100μm. (C) Representative image of 24hpi orthotopic injection of zebrafish at 72hpf (top) with an inset (bottom). Micrometer bar: 100μm. (D) GSCs, HF3077 (left) and HF3160 (right) injected into yolk sac. Tumour burden scored at 24hpi using ImageJ. Representative images (top) with insets (bottom). Signal quantified as Integrated Density Values (IDV). Micrometer bar: 750μm. (E) Fluorescent images of GSC HF3077-derived tumour foci growth in zebrafish embryos 72 hrs, 96 hrs, & 5 days post injection, quantified as number (no.) of tumour foci (left) and intensity of the fluorescent signal (right) as IDV using ImageJ software. Micrometer bar: 300 μm. HF3077 cells labeled with DiI.

### Quantifying proximal vs. distal migration in zebrafish PDXs

GSCs HF3077 (DiO, green) and HF3160 (DiI, red) were injected into the yolk sac at 72hpf (Fig. 2A). Cell migration was scored at 72hpi (Fig. 2B). Interestingly, only HF3160, mesenchymal GSCs, demonstrate migration and invasion across zebrafish larvae tested at 48hpi and the number of proximal foci established by HF3160 is decreased in comparison to distal foci (Fig. 2B). No migrated GSCs of HF3077 were detected at 48hpi. HF3077 cells demonstrated delayed migration in comparison to HF3160 GSCs with proximal and distal foci observed 4dpi (Fig. 2C).

**Figure 2.**
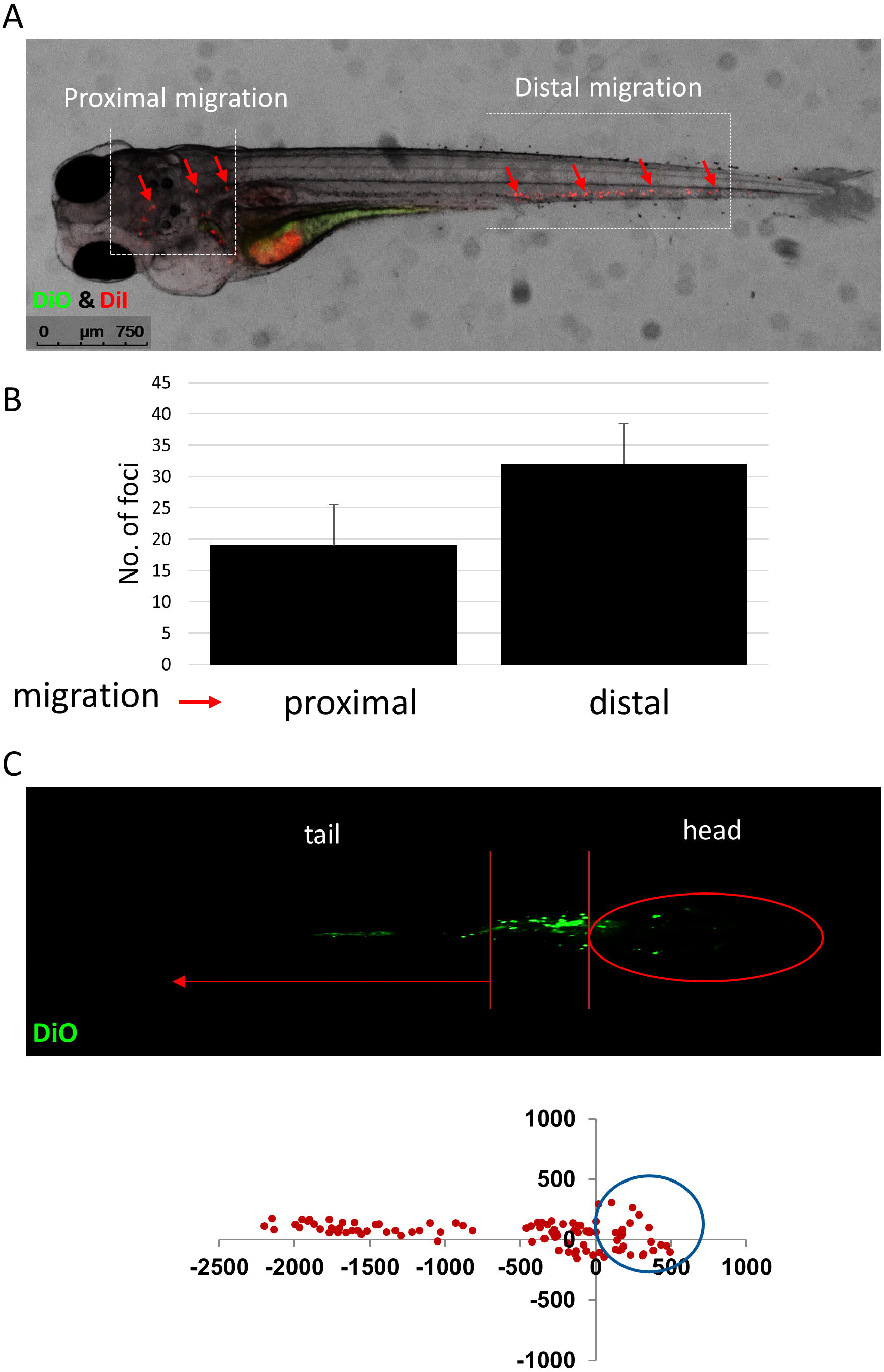
Directionality and migration of tumour cells from the site of injection in zebrafish embryos injected into the yolk sac with GSCs HF3077 (DiO, green) and HF3160 (DiI, red). (A) Migration of tumour cells in proximity to the injection site (proximal metastasis) and cells forming distal metastasis (B) quantification of migrated cells. (C) Graphs representing coordinates of each tumour foci of HF3077 detected at 4dpi with 0,0 being the injection site.

### Zebrafish transgenic models of tumour microenvironment

Several zebrafish models are available to study diverse components of the tumour microenvironment including endothelial cells and vasculature using Tg(fli1a:dsRED) or cells of the immune system using Tg(mfap4.1:mCherry, tnfamodeGFP) and Tg(mpx:EGFP) (Fig. 3A) transgenic zebrafish. Tg(fli1a:dsRED) zebrafish were injected with HF3077 GSCs labeled with DiO and small number of cell was detected in vasculature of the brain at 72hpi (Fig. 3B). Same model used for orthotopic injection of HF3077 GSCs labeled with DiD tracker demonstrated improved capacity to visualize GSC interactions with endothelial cells of the brain at 48hpi (Fig. 3C).

**Figure 3.**
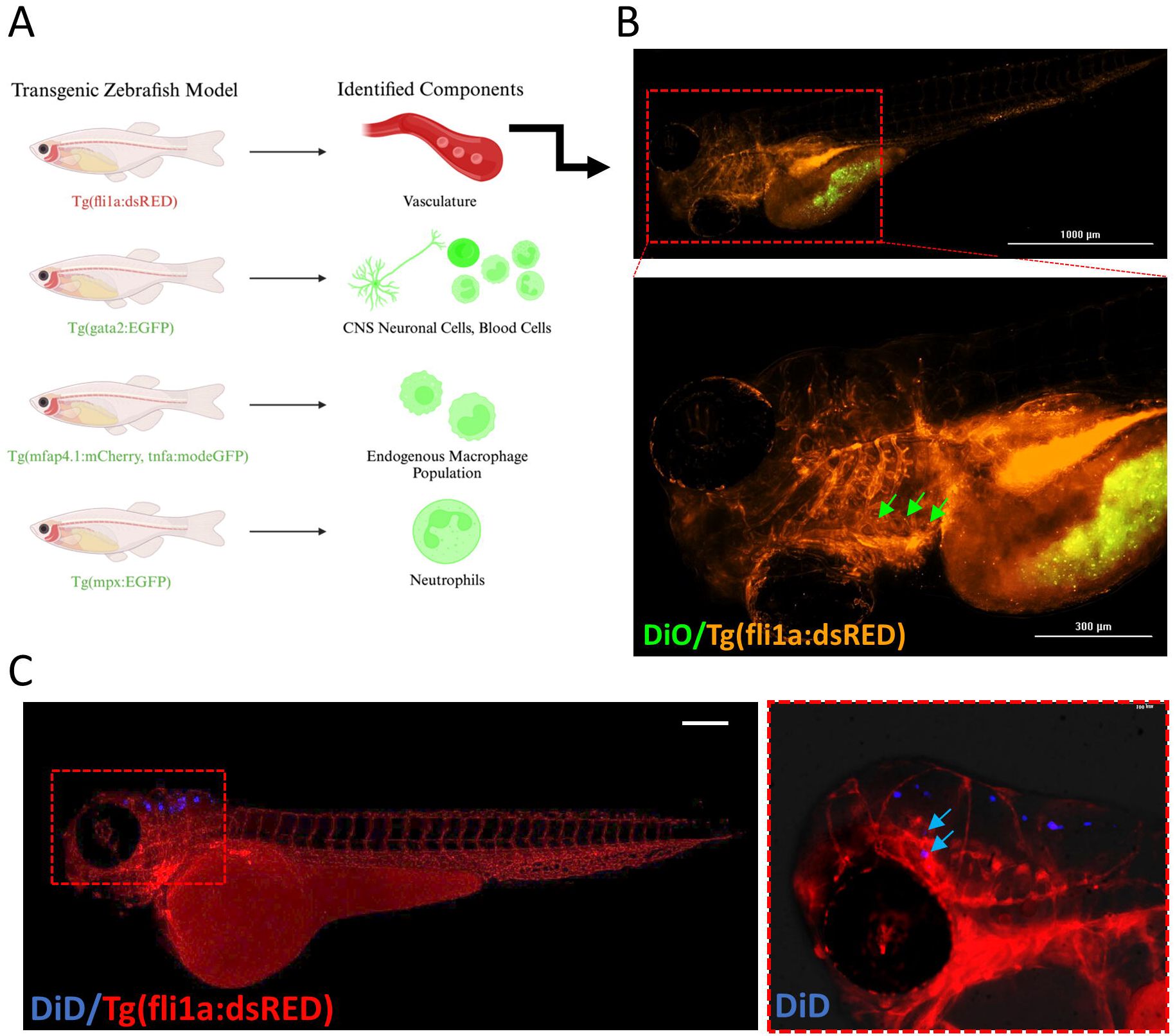
Zebrafish models suitable for studies of tumour microenvironment in glioma. (A) Schema presenting commercially available models. Created with BioRender.com. (B) Representative 3D image of Tg(fli1a:dsRED) zebrafish embryos at 72hpf injected with GSCs HF3077 into the yolk sac at 72hpi. Images obtained with Biotek Cytation 5 Live Cell Imager RFP filter. (C) Orthotopic injection at 72hpf with GSCs HF3077 labeled with DiD into Tg(fli1a:dsRed) zebrafish embryos; image (left) with inset (right) taken at 48hpi. Micrometer bar: 200 μm. Images obtained with Leica Stereo Microscope Cy3 filter.

### Detecting tumour foci post GSC injection in zebrafish PDXs

Zebrafish PDXs are collected processed, sectioned, and stained as shown in Fig. 4A. Upon examination of the sections stained with H&E of the entire zebrafish larvae (Fig. 4B) and regions of the head and brain (Fig. 4C), tumour foci previously observed in Fig. 1-3 could not be detected using standard H&E staining. The sections were then stained with an antibody against PD-L1 protein allowing for specific detection of tumour cells (Fig. 4D, red arrows). Lymphocyte recruitment around the tumour cells was detected using positive an antibody against LCP expression (green arrows).

**Figure 4.**
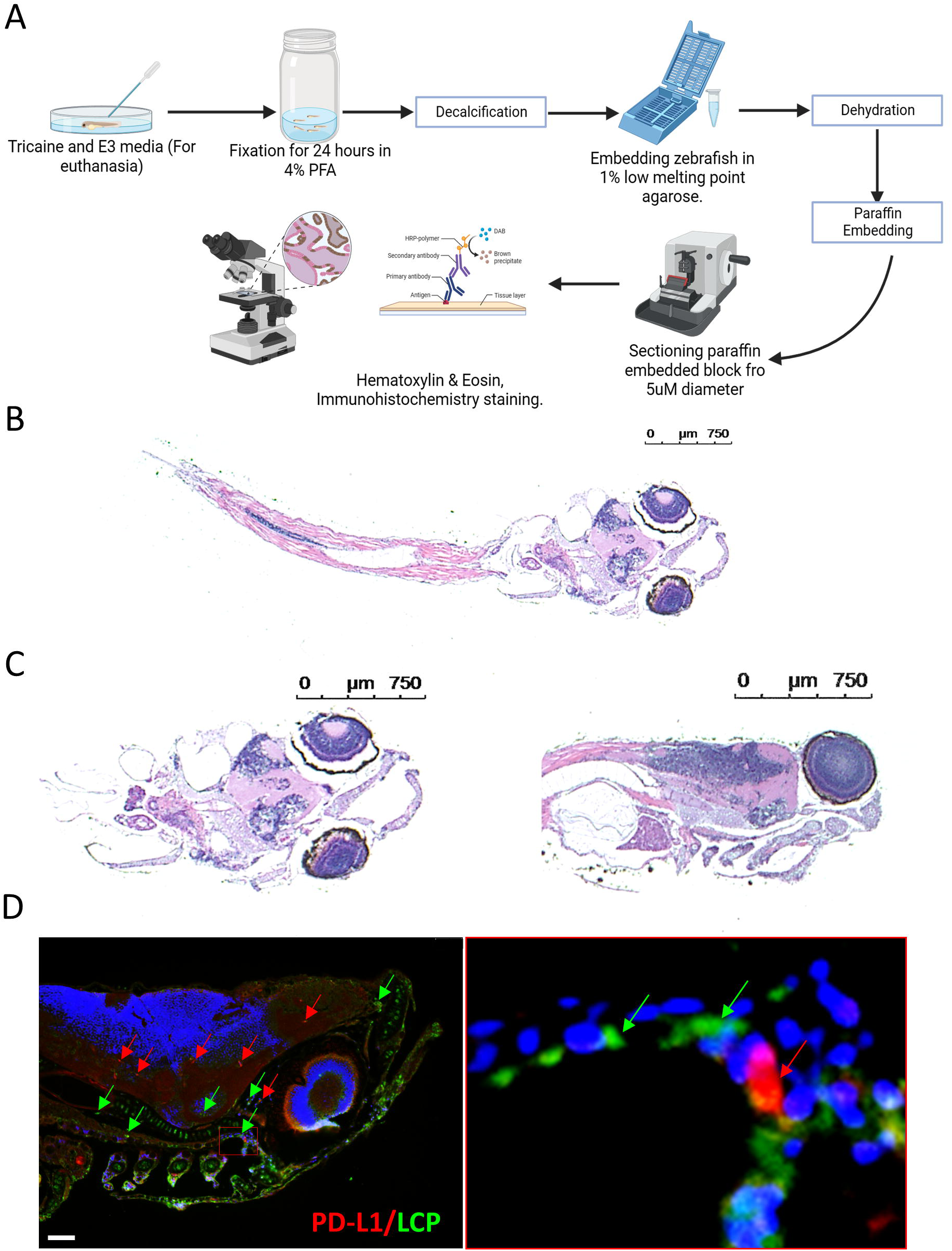
(A) Schematic representation of zebrafish embryo processing for sectioning and embedding. Created with BioRender.com. (B) H&E staining of longitudinal cross sections of zebrafish embryos; representative images. (C) Representative H&E images of zebrafish head/brain region at 10dpf stained with H&E. (D) Fluorescent image (left) with inset (right) of sections obtained from zebrafish PDX model at 6 days post injection stained for Programmed Death Ligand 1 (PD-L1) found on injected cells; red arrows, and Lymphocyte Cytosolic Protein (LCP); green arrows. Micrometer bar: 40μm.

## References

1. Angom RS, Nakka NMR, Bhattacharya S. Advances in Glioblastoma Therapy: An Update on Current Approaches. Brain Sci. 2023;13(11):1536. doi:10.3390/brainsci13111536

2. Rong L, Li N, Zhang Z. Emerging therapies for glioblastoma: current state and future directions. J Exp Clin Cancer Res. 2022;41(1):142. doi:10.1186/s13046-022-02349-7

3. Song Q, Ruiz J, Xing F, et al. Single-cell sequencing reveals the landscape of the human brain metastatic microenvironment. Commun Biol. 2023;6(1):1–13. doi:10.1038/s42003-023-05124-2

4. Liu Z, Wang S, Yu K, et al. The promoting effect and mechanism of MAD2L2 on stemness maintenance and malignant progression in glioma. J Transl Med. 2023;21:863. doi:10.1186/s12967-023-04740-0

5. Tang X, Zuo C, Fang P, Liu G, Qiu Y, Huang Y, Tang R. Targeting Glioblastoma Stem Cells: A Review on Biomarkers, Signal Pathways and Targeted Therapy. Front Oncol. 2021;11:701291. doi:10.3389/fonc.2021.701291

6. Glumac PM, LeBeau AM. The role of CD133 in cancer: a concise review. Clin Transl Med. 2018;7(1):18. doi:10.1186/s40169-018-0198-1

7. Hassn Mesrati M, Syafruddin SE, Mohtar MA, Syahir A. CD44: A Multifunctional Mediator of Cancer Progression. Biomolecules. 2021;11(12):1850. doi: 10.3390/biom11121850

8. de Visser KE, Joyce JA. The evolving tumour microenvironment: From cancer initiation to metastatic outgrowth. Cancer Cell. 2023;41(3):374–403. doi:10.1016/j.ccell.2023.02.016

9. Nowosad A, Marine JC, Karras P. Perivascular niches: critical hubs in cancer evolution. Trends in Cancer. 2023;9(11):897–910. doi:10.1016/j.trecan.2023.06.010

10. Zhou Y, Xia J, Xu S, et al. Experimental mouse models for translational human cancer research. Front. Immunol. 2023;14:1095388. doi:10.3389/fimmu.2023.1095388

11. Ali Z, Vildevall M, Rodriguez GV, et al. Zebrafish patient-derived xenograft models predict lymph node involvement and treatment outcome in non-small cell lung cancer. J Exp Clin Cancer Res. 2022;41(1):58. doi:10.1186/s13046-022-02280-x

12. Ji X, Chen S, Guo Y, et al. Establishment and evaluation of four different types of patient-derived xenograft models. Cancer Cell Int. 2017;17:122. doi:10.1186/s12935-017-0497-4

13. Pudelko L, Edwards S, Balan M, et al. An orthotopic glioblastoma animal model suitable for high-throughput screenings. Neuro-Oncol. 2018;20(11):1475–1484. doi:10.1093/neuonc/noy071

14. Choi TY, Cho TI, Lee YR, et al. Zebrafish as an animal model for biomedical research. Exp Mol Med. 2023;53:310–317. doi:10.1038/s12276-021-00571-5

15. Copper JE, Budgeon LR, Foutz CA, et al. Comparative analysis of fixation and embedding techniques for optimized histological preparation of zebrafish. Comp Biochem Physiol X Toxicol Pharmacol. 2018;208:38–46. doi:10.1016/j.cbpc.2017.11.003

